# The peach RGF/GLV signalling peptide pCTG134 is involved in a regulatory circuit that sustains auxin and ethylene actions

**DOI:** 10.1101/141705

**Authors:** Nicola Busatto, Umberto Salvagnin, Francesca Resentini, Silvia Quaresimin, Lorella Navazio, Oriano Marin, Maria Pellegrini, Fabrizio Costa, Dale F. Mierke, Livio Trainotti

## Abstract

Peach is a climacteric species whose ripening is regulated by the plant hormone ethylene. A crosstalk mechanism with auxin is necessary to support climacteric ethylene synthesis. The homeostasis control of auxin is regulated also by the activity of peptide hormones (PHs), acting both as short and long distant ligands. In this work, we investigated the role of *CTG134*, a peach gene encoding a GOLVEN-like PH isolated in mesocarp at the onset of ripening.

In peach fruit, *CTG134* was expressed during the climacteric transition and its mRNA level was induced by auxin and 1-methylcyclopropene (1-MCP) treatments, whereas it was minimally affected by ethylene. To better elucidate its function, *CTG134* was overexpressed in *Arabidopsis* and tobacco, which showed abnormal root hair growth, similar to wild-type plants treated with a synthetic form of the peptide. Molecular surveys demonstrated an impaired hormonal crosstalk, resulting in a re-modulated expression of a set of genes involved in both ethylene and auxin domains. In addition, the promoter of pCTG134 fused with GUS reporter highlighted gene activity in plant organs in which the auxin-ethylene interplay is known to occur. These data support the role of pCTG134 as mediator in an auxin-ethylene regulatory circuit.

**Highlight:** The role of the peach RGF/GLV peptide during root hair formation in *Arabidopsis* and tobacco supports its involvement in a cross-hormonal auxin-ethylene regulatory circuit.

## Introduction

In Angiosperms, fruits, besides providing essential and beneficial compounds to the human diet, protect the seeds enabling their dispersion at the end of the ripening phase. These organs originally develop from the ovary, with the possible contribution of other flower parts. According to the physiological regulation of ripening, fleshy fruits can be distinguished into climacteric (such as tomato, peach and apple) and non climacteric (such as strawberry, grape and citrus), depending on the presence of a burst in the production of the plant hormone ethylene accompanied by a respiratory increase occurring at the late stage of fruit ripening (Liu *et al*., 2015). Ethylene is produced through the sequential activation of two biosynthetic systems (McMurchie *et al*., 1972; Oetiker *et al*., 1997). The auto-inhibitory system 1, found in both climacteric and non-climacteric fruit, maintains a basal level of ethylene during the vegetative growth of plants as well as in wound and stress response. The autocatalytic system 2 produces, instead, a much larger amount of the hormone, typical of climacteric fruits in full-ripening phase. In the tomato model, the switch between the two systems is based upon the differential expression of 1-amino-cyclopropane-1-carboxylic acid (ACC) synthase (ACS) and ACC oxidase (ACO) genes during the ripening process (Barry *et al*., 2000). The transition from the system 1 to system 2, which represents a crucial point in the ripening process of climacteric species (Cara and Giovannoni, 2008), is also regulated by genetic (Vrebalov *et al*., 2002; Manning *et al*., 2006) and epigenetic factors (Zhong *et al*., 2013), together with the surrounding environmental effect and interplay with other plant hormones (Klee and Giovannoni, 2011).

Emerging evidence suggests that relative functions of plant hormones are not restricted to a particular stage only. A complex network of more than one plant hormone is therefore involved in controlling various aspects of fruit development (Kumar *et al*., 2014). The knowledge of the hormonal network and crosstalk relationship between different hormones during the stages of the fruit life cycle is still far from being complete (Kumar *et al*., 2014) and relies almost entirely on model species (*Arabidopsis* and tomato). The action of the phytohormones depends not only on the cellular context, but also on the relationship established among different hormones. To date, the hormonal crosstalk has been mainly investigated in *Arabidopsis*, which shed light, among others, on the crosstalk between auxin and ethylene (Poel *et al*., 2015). The first and most evident effect of the interaction between these two hormones is on the regulation of the root morphogenesis process. Indeed, in this organ it has been demonstrated that root hair formation, elongation (Pitts *et al*., 1998; Dolan, 2001) and differentiation as well as the development of lateral roots are regulated by the interplay occurring between auxin and ethylene (Zhang *et al*., 2016). On the other hand, ethylene can also modify the auxin patterning by modulating IAA transport (Prayitno *et al*., 2006). Cellular and genetic evidences have shown a physiological connection between hormones and peptide hormones (PHs). ROOT GROWTH FACTOR/GOLVEN/CLE-Like (RGF/GLV/CLEL) peptides can in fact alter auxin gradients by changing the turnover of IAA carriers (Whitford *et al*., 2012). Despite the importance of this regulatory mechanism, the biology of PHs is still in its infancy, especially in non-model but agronomically relevant species.

Auxin and ethylene have been described to interact at the level of ethylene biosynthesis (Abel *et al*., 1995) not only in *Arabidopsis* roots but also during the ripening of different fruit species, such as tomato (Abel *et al*., 1996), peach (Trainotti *et al*., 2007) and apple (Shin *et al*., 2015). Although the molecular mechanisms of the interplay between auxin and ethylene during fruit ripening are still unknown, recent data suggest that PHs (Matsubayashi, 2014; Tavormina *et al*., 2015) could be the crossroads between the two hormones in peach (Tadiello *et al*., 2016). One PH in particular, namely CTG134 GLV-like, was identified through a comprehensive survey carried out with the μPeach1.0 (Tadiello et al. 2016). This gene was expressed at the transition step between the preclimacteric and the climacteric stage. Moreover, while CTG134 was induced by exogenous treatment of 1-methylcyclopropene (1-MCP), an ethylene competitor largely used to delay the normal physiological ripening progression (Watkins, 2006), its expression was also totally repressed in ripe fruit of *stony hard*, a peach mutant showing impairment both in ethylene production and cell wall metabolism (Pan *et al*., 2015).

In this work the peptide pCTG134, isolated from peach, was functionally validated in *Arabidopsis* and tobacco, providing new evidence about its role as a major regulator in the auxin-ethylene crosstalk.

## Materials and methods

### Plant materials

Peach fruit s were collected from cv. ‘Redhaven’ (RH) as described in Tadiello *et al.* (2016). The heterologous CTG134 overexpression was carried out in *Arabidopsis* and tobacco plants. Seeds of *Arabidopsis thaliana* Columbia accession (Col-0) were surface-sterilized, stratified overnight at 4°C and germinated on plant growth medium (Murashige and Skoog, 1962) or in potting soil at 22°C. To characterize root growth, MS plates were tilted with an angle of 45°. *Nicotiana tabacum* SNN plants were instead grown following standard protocols in controlled greenhouse.

### Hormone treatments on Redhaven fruit

The auxin treatment was performed by dipping the whole fruit in 1-naphthalene acetic acid [NAA, 2 mmol L^−1^ added with Silwet L-77 (200 μL L^−1^) as surfactant] for 15 min; thereafter, fruit were sprayed with the NAA solution every 12 h over a period of 48 h (NAA omitted in the mock control). The ethylene treatment was instead carried out as previously described in (Tadiello *et al*., 2016).

### 1-MCP treatments on Stark Red Gold fruit

Treatment of cv. “Stark Red Gold” (SRG) peach fruits with 1-MCP was carried out as described in Tadiello *et al.* (2016).

### RNA extraction and expression analyses by quantitative Real time PCR (qRT-PCR)

Peach RNA was prepared from a frozen powder obtained by grinding mesocarp sectors from at least four different fruits. From four grams of this powder, total RNA was extracted following a protocol previously described (Chang *et al*., 1993). *Arabidopsis* RNA was extracted from wild type and 35S::CTG134 mutant seedlings, using the LiCl method (Verwoerd *et al*., 1989). Expression analyses were performed using Power SYBR Green PCR Master Mix (Applied Biosystems). Normalization was performed using UBIQUITIN10 (UBI10) and ACTIN8 as internal standards for Arabidopsis and Ppa009483m/Prupe.8G137600 for peach (Primers are listed in Table S1). qRT-PCR was performed and the obtained data analysed as previously described (Tadiello *et al*., 2016).

### In-situ hybridizations

*Prunus persica* S3II and S4 fruits were fixed and embedded in 4% paraformaldehyde. A *CTG134* specific probe was amplified by PCR from S3II and S4 fruit cDNAs (primers listed in Table S1) and further cloned in pGEM T-easy vector (Promega). The CTG134 transformed vector was further used as template for the creation of sense and antisense probes by an *in-vitro* transcription performed with SP6 and T7 polymerases. Sections of plant tissue were probed with dioxigenin-labelled antisense RNA-probe as previously described (Brambilla *et al*., 2007) and observed with a Zeiss Axiophot D1 light microscope (http://www.zeiss.com).

### pPR97-proCTG134:GUS construct design

To assess the CTG134 promoter activity, a fragment of 2679 bp located upstream of the coding sequence initiation site (Supplementary Fig. S1A) was isolated from peach genomic DNA (cv Red Haven) by PCR. PCR product was cloned into the pCR8/GW/TOPO TA Cloning vector (Invitrogen, Carlsbad, CA, USA), according to the manufacturer’s instructions and confirmed by sequencing. The promoter fragment was thus subcloned into a pPR97-derived vector (12.20 kb), made compatible with the Gateway cloning system (LR Clonase II – Invitrogen, Carlsbad, CA, USA). This modified pPR97 vector with kanamycin resistance was employed for stable transformations both in *Arabidopsis thaliana* and *Nicotiana tabacum*, to measure the CTG134 promoter activity. The promoter was tested by cloning the upstream sequence and a GUS reporter gene interrupted by a plant intron (Vancanneyt *et al*., 1990). To make easier the cloning, a CC_rfA gateway cassette was inserted (SmaI) upstream of the reporter gene and the antibiotic kanamycin was used to select resistant successfully transformed plants.

### pGreen-AmpR-KanNos-35S:CTG134 construct design

The CTG134 coding sequence (524 bp) was amplified by PCR from *Prunus persica* (cv Red Haven, S4I development stage) cDNA and subsequently cloned into the pCR8/GW/TOPO TA Cloning vector (Invitrogen, Carlsbad, CA, USA). The CTG134 CDS was further inserted into a pGreen-derived vector (Hellens *et al*., 2000) with the Gateway cloning system (LR Clonase II – Invitrogen, Carlsbad, CA, USA). The pGreen-derived vector was modified to confer resistance to both kanamycin and ampicillin. Moreover, a CC_rfA gateway cassette was inserted downstream of the 35S promoter in the EcoRV site. As before, the selection of plants was carried out with kanamycin (Supplementary Fig. S1B).

### Arabidopsis thaliana and tobacco transformation

Single PCR-positive *Agrobacterium* GV3101 colonies were used to grow liquid cultures for the transformation of *Arabidopsis thaliana* Columbia 0 plants with the floral dip method (Clough and Bent, 1998). The first flowers of four-weeks old plants were cut to allow, after 4-8 days, the growing of a second set, further dipped in a suspension of *Agrobacterium* cells (OD600 = 0.8), sucrose (5% m/v) and Silwet L-77 (0.05%). Plants were incubated in the dark for 16 hours before a second growing phase in growth chamber (16/8 light/dark cycle, 25°C, 70% relative humidity) until seeds were obtained. Transformed plants were screened on solid ½ MS medium (MS salts with vitamins 2.17 g/L, sucrose 15 g/L, pH 5.75) supplemented with kanamycin (50 mg L^−1^). After one week, the resistant plants were planted into soil and grown in greenhouse for at least two generations, until T-DNA insertions reached homozygosity. Plants were screened for the presence of the transgene by PCR on genomic DNA using specific primer pairs.

In-vitro grown *N. tabacum* SNN plants were instead transformed following the protocol reported by (Fisher and Guiltinan, 1995). As for Arabidopsis, plants were screened for the presence of the transgene with PCR on genomic DNA using specific primer pairs.

### Peptide synthesis

The peptides DYSPARRKPPIHN and DY(SO_3_H_2_)SPARRKPPIHN were synthesized by automatic solid phase procedures. The synthesis was performed using a multiple peptide synthesizer (SyroII, MultiSynTech GmbH) on a pre-loaded Wang resin (100-200 mesh) with N-α-Fmoc-N-β-trityl-l-asparagine (Novabiochem, Bad Soden, Germany). The fluoren-9-ylmethoxycarbonyl (Fmoc) strategy (Fields and Noble, 1990) was used throughout the peptide chain assembly, utilizing O-(7-azabenzotriazol-1-yl)-N,N,N′,N′-tetramethyluronium hexafluorophosphate (HATU) as coupling reagent (Carpino *et al*., 2001). The side-chain protected amino acid building blocks used were: N-α-Fmoc-β-tert-butyl-l-aspartic acid, N-α-Fmoc-Nε-tert-butyloxycarbonyl-l-lysine, N-α-Fmoc-Nω-2,2,4,6,7-pentamethyldihydrobenzofuran-5-sulfonyl-l-arginine, N-α-Fmoc-O-tert-butyl-l-serine, N-α-Fmoc-N(im)-trityl-l-histidine, N-α-Fmoc-O-tert-butyl-l-tyrosine and N-α-Fmoc-O-sulfo-l-tyrosine tetrabutylammonium salt. Cleavage of the peptides was performed by incubating the peptidyl resins with trifluoroacetic acid/H2O/triisopropylsilane (95%/2,5% /2,5%) for 2.5 h at 0 °C. Crude peptides were purified by a preparative reverse phase HPLC. Molecular masses of the peptide were confirmed by mass spectroscopy on a MALDI TOF-TOF using a Applied Biosystems 4800 mass spectrometer.

### Ca^2+^ measurement assays

Ca^2+^ measurement assays were carried out in *Arabidopsis* cell suspension cultures obtained from *Arabidopsis* seedlings stably expressing cytosolic aequorin (seeds kindly provided by M.R. Knight, Durham, UK). Reconstitution of aequorin and Ca^2+^ measurements were carried out as described (Sello *et al*., 2016).

## Results

### Regulation of CTG134 expression

Expression of *CTG134* was assessed in peach mesocarp during the onset of fruit ripening (i.e. at early stage 4 – S4I – Fig. 1A). CTG134 mRNA accumulated in preclimacteric fruit (i.e. S3II) after auxin treatment, while exogenous ethylene had no effect (Fig. 1B). Moreover, treatment with the ethylene inhibitor 1-MCP induced *CTG134* transcription at stages before (cl 0) and coincident (cl 1) with the full climacteric (Fig. 1C). The peach mesocarp at ripening is mainly made up of parenchymal cells and vascular tissue (Zanchin *et al*., 1994). To localize the types of cells expressing *CTG134* at ripening, *in-situ* hybridization experiments were carried out with mesocarp sections prepared by peach fruit in S4 stage. The CTG134 mRNA was localized in vascular bundles (Supplementary Fig. S2C), most likely in the phloem or parenchymal cells (Fig. 1D).

**Fig 1.**
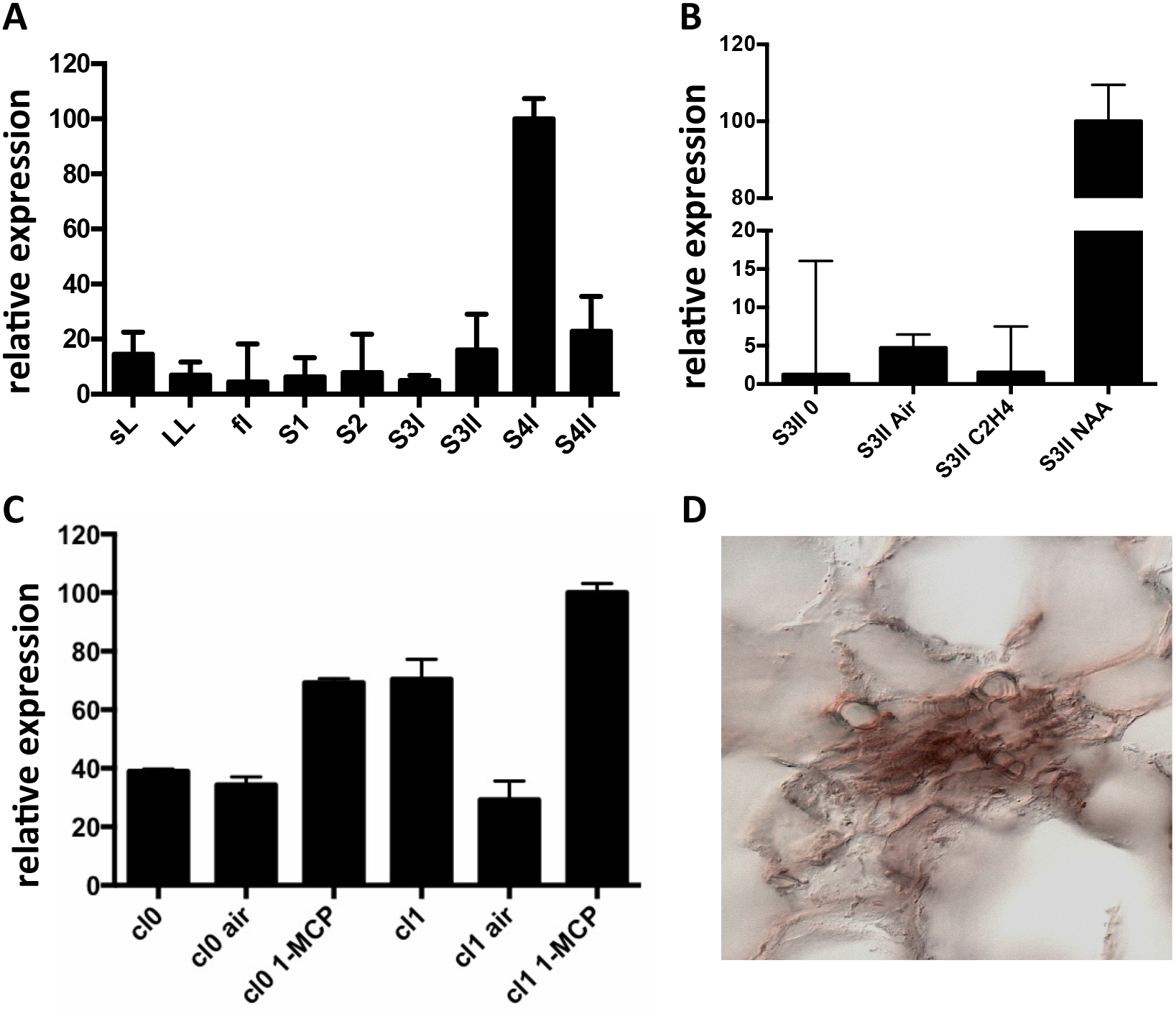
Expression profile of *CTG134*. (A) *CTG134* expression was barely detectable, by qRT-PCR, in non-fruit organs (small expanding leaves-sL- and fully developed leaves –LL-) and in fruit at early development (stage 1 and 2, - S1, S2-). In mature fruit (S3II) there was a sharp increase in *CTG134* transcription (S4I), slightly ceasing after the ethylene peak (S4II). (B) Ethylene, auxin and 1-MCP responsiveness of *CTG134. CTG134* expression, barely detectable in mature preclimacteric fruit (S3II 0) was strongly increased upon auxin (NAA, 1-naphthalene acetic acid, a synthetic auxin) but not ethylene treatment. (C) Both in Class 0 and Class 1 S4 fruit, 1-MCP upregulated *CGT134* expression. (D) Localization of *CTG134* expression in peach mesocarp by *in-situ* hybridization. In peach mesocarp at S4, *CTG134* expression was mainly associated with vascular bundles (control sections in Fig. S2). Scale bar = 50 μm.

Since peach is a recalcitrant species to transform, *proCTG134:GUS* lines were generated in both tobacco and *Arabidopsis* model species. In tobacco, a slight but evident GUS staining was detected in the apical meristem (RAM) of *in-vitro* grown lateral roots (Fig. 2A). Moreover, a dark staining was visible in lateral root emergence (Fig. 2B) as well as in leaf, mainly associated, but not limited to, the vascular tissue (Fig. 2C). In the stem of one-week-old plantlets, GUS expression was localized in phloem of cell layers closed to the cambium (Fig. 2D). GUS expression was also tested in reproductive organs, where it was detected in the tips of both young sepals and petals (not shown) and in capsules at the level of the dehiscence zone (Fig. 2E). The inner part of the fruit was the part more significantly stained (Figures 2F and 2G), with the highest expression in the placenta (Fig. 2G). On the contrary, in all the transgenic lines investigated in this study, the GUS colouration was never observed in ovule. In one-week-old tobacco seedlings the reporter was more expressed in cotyledons than roots. However, five-hour treatment with 50μM IAA induced a different GUS staining in the entire shoot apex and root, reaching the highest intensity in the root-stem transition zone (Fig. 2H). A similar auxin-induced expression was also observed in roots of *in-vitro* grown plantlets (Fig. 2I). The stimulation of the GUS staining in tobacco finds also consistency with the aforementioned expression pattern of CTG134 in peach fruit. The expression of this element was in fact enhanced by auxin (Fig. 1B) and auxin responsive elements (AREs) were moreover detected in the *CTG134* promoter region (Supplementary Fig. S1B). To further validate the heterologous analysis carried out in tobacco, the activity of the CTG134 promoter was additionally investigated in *Arabidopsis* (Fig. 3A). Also in this species, the GUS expression was higher in cotyledons (Fig. 3B) rather than in primary root, where the GUS staining was undetectable in the RAM (Fig. 3C). The GUS activity was instead clearly visible at the root-stem transition zone (Fig. 3D) and during lateral root emergence (Fig. 3E). In the reproductive organs, the expression pattern was detected in abscission zones before (Fig. 3F) and after (Fig. 3G) shedding. The expression was also detected in maturing siliques and leaves, especially in those associated with vascular bundles (Fig. 3H).

**Fig 2.**
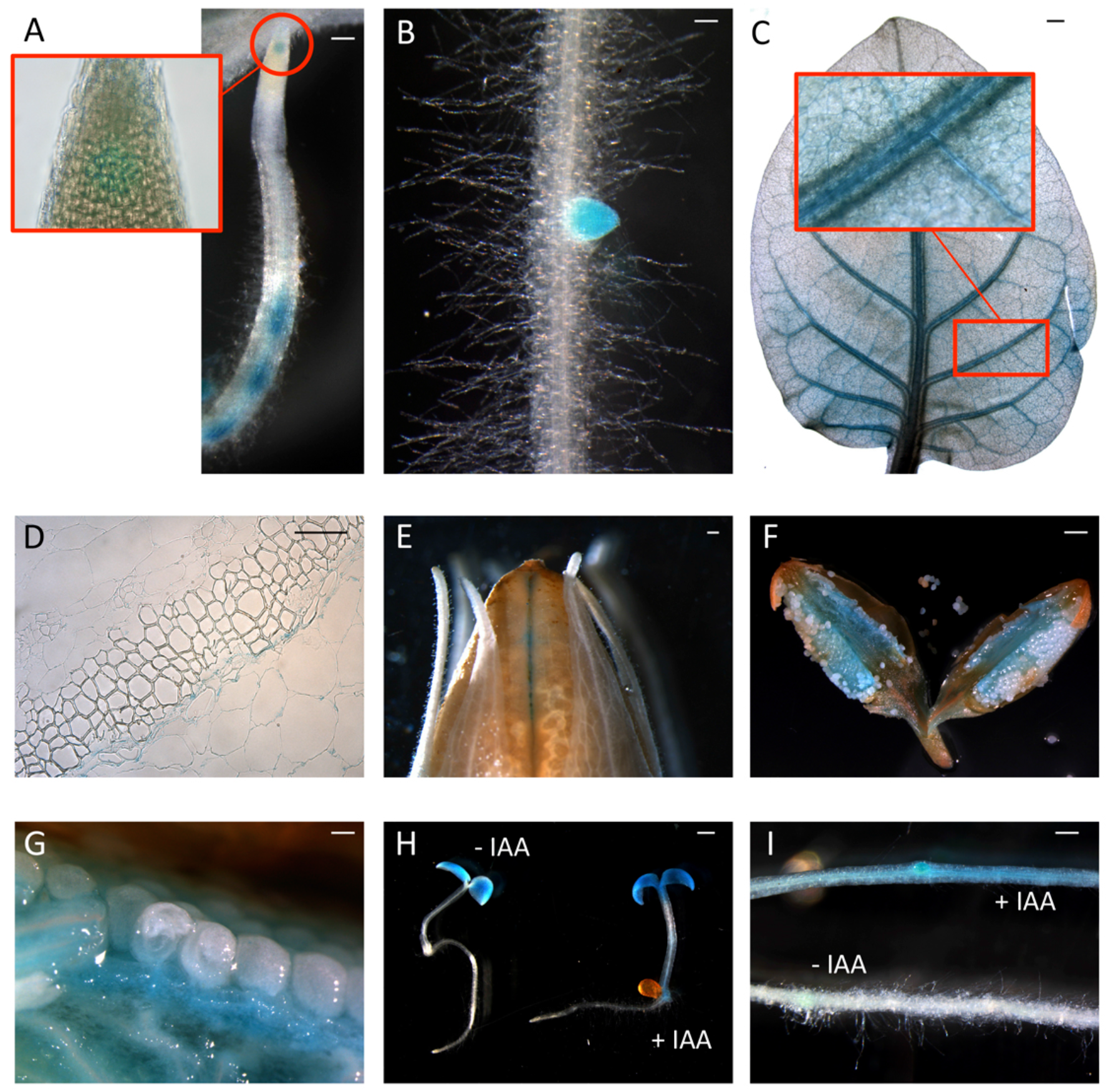
*ProCTG134:GUS* expression in tobacco and auxin responsiveness. (A, B) In tobacco roots the expression of GUS was detected at the level of the RAM (inset) but mainly at the level of lateral root primordia. (C) Staining was detectable also in leaves, especially if treated with 50μM IAA, and particularly in veins (inset). (D) In the stem, GUS expression was more abundant in parenchymatic cells of the vascular tissue. In the fruit expression was visible at the dehiscence zone (E) and in the placenta (F, G). (H) Auxin responsiveness in one-week-old representative seedlings (untreated on the left, and treaded with 50μM IAA on the right) and in the root (I, untreated, on the bottom, and treaded with 50μM IAA, on the top). Scale bar in the panels B, C and F = 500 μm, in A = 200 μm, in D = 100 μm and in E = 1000 μm.

**Fig 3.**
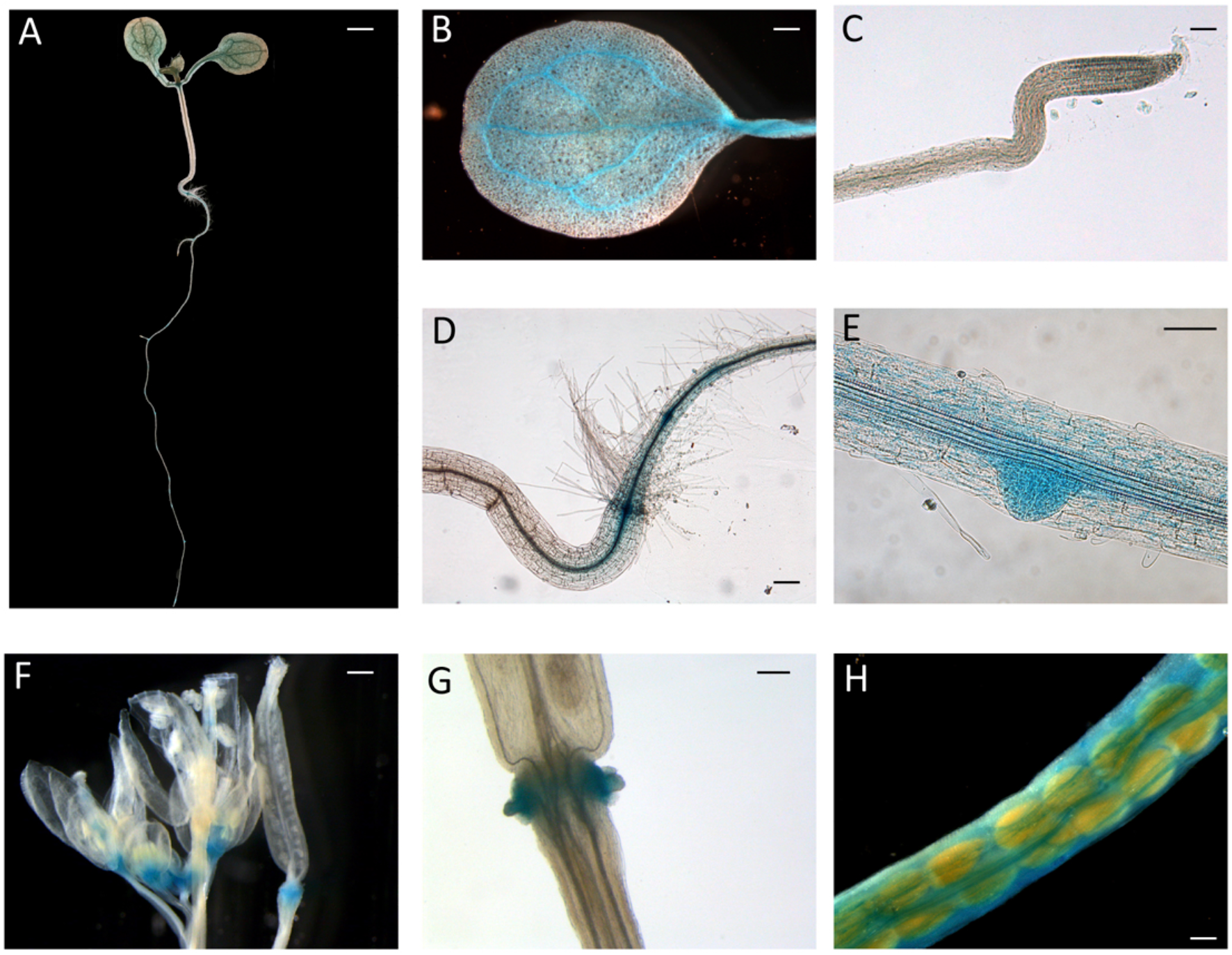
*ProCTG134:GUS* expression in *Arabidopsis.* At seven days after germination (A), GUS staining is detectable in cotyledons, especially in veins (B), at the root-shoot transition zone (D) and in lateral root primordia (E), while is barely detectable in RAM (C). In the reproductive part, expression was detected in abscission zones before (F) and after (G) organ shedding. Expression was detectable also in maturing siliques mainly associated with vascular bundles (H). Scale bar in the panels B, C and F = 500 μm, in A = 200 μm, in D = 100 μm and in E = 1000 μm.

### 4.2 Hormonal regulation of CTG134 in tobacco

To test whether the auxin responsiveness was due to the promoter regulatory region, one-week old tobacco seedlings of line #2 were exposed to increasing concentrations of IAA. The CTG134 promoter was responsive to IAA already at 0.5μM, with an activity pattern proportional to the hormone concentrations. The system reached saturation at 50μM (Fig. 4A). The IAA induction kinetic was assessed over a time course of 20 hours on tobacco seedlings of line #2 treated with 10μM IAA. An initial slight induction in both control and treated samples was observed already after 30 minutes, after which the GUS activity remained at a basal level in the control, while in the IAA treated samples a significant burst was observed after 3 hours after the treatment (Fig. 4B). Since in peach fruit the expression of CTG134 was insensitive to ethylene and induced by 1-MCP (Fig. 2B and 2C), the promoter responsiveness was tested by treating ten-day-old tobacco seedlings for sixteen hours with ethylene (10μL L^−1^), IAA (10μM) and 1-MCP (1μL L^−1^). 1-MCP induced the reporter activity similarly to auxin (Fig. 4C), while treatment with ethylene did not change the expression of the GUS reporter gene.

**Fig 4.**
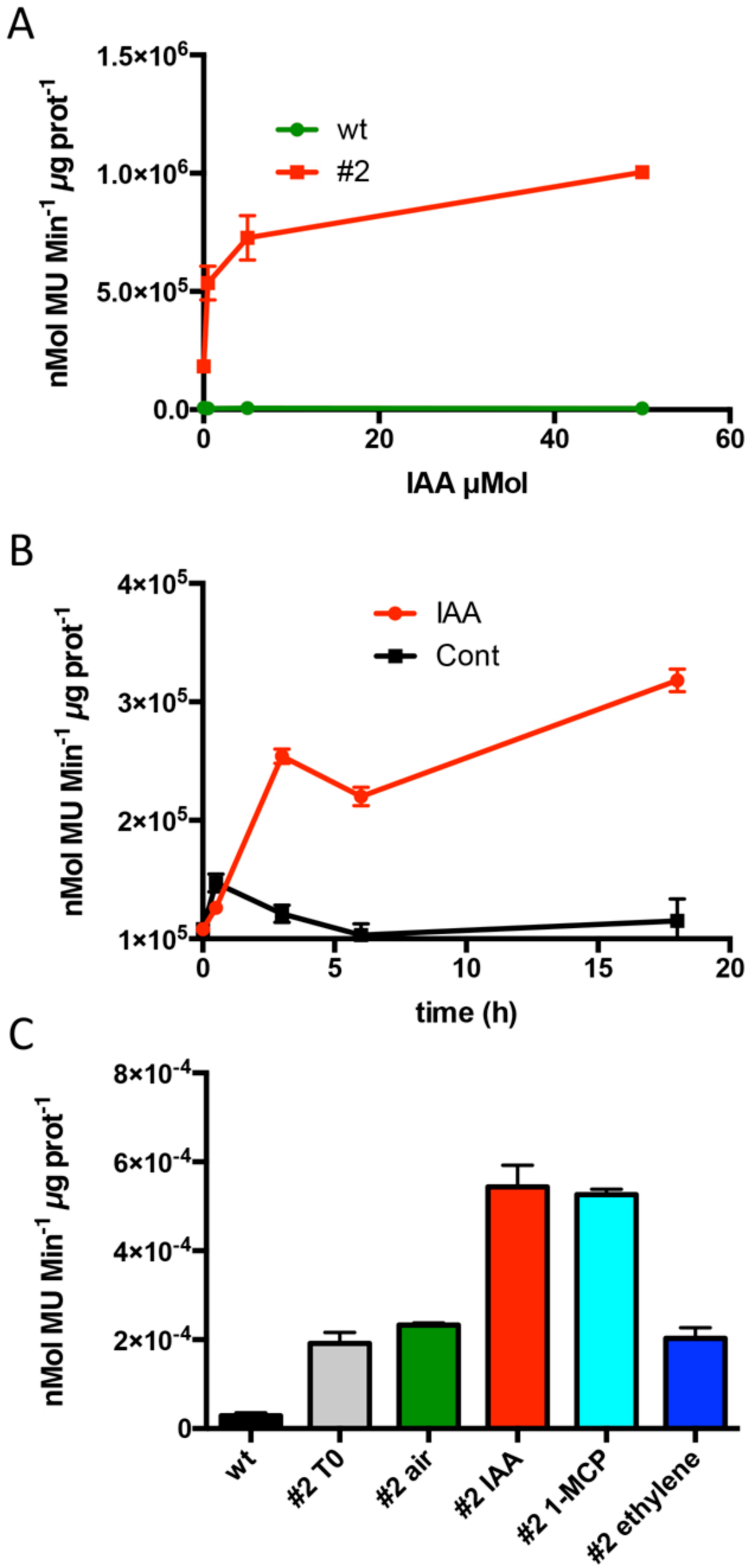
Hormone responsiveness of *ProCTG134:GUS* in tobacco seedlings. (A) Auxin was effective in inducing the promoter of *CTG134* already at 0.5 μM, to reach almost complete saturation at 50 μM. (B) Saturation of the auxin induction after three hours. (C) Besides auxin, also 1-MCP had an inductive effect on the promoter of *CTG134*, while ethylene seemed ineffective. All experiments were carried out with T3 seedlings of line #2.

### *Over-expression of* CTG134 *in tobacco*

To functionally investigate the role of the peptide CTG134 peptide, its full-length coding sequence, under the control of the Cauliflower mosaic virus 35S (35S CaMV) promoter was expressed in tobacco. The development of longer root hairs was noticed already in the early phases of transgenic plant production (Fig. 5A). A YFP gene, cloned in the same binary vector as CTG134, was overexpressed to have control plants able to grow on kanamycin and gentamicin present in the growth media. To further assess this phenotype, scions from different clones were propagated and primary roots from 30 day-old plants were analysed by taking images in the root portion located at 6 mm from the root tip. On average, the CTG134 overexpressing lines showed an increase of at least two-fold in root hair length (ANOVA, F = 87.75, df = 155, p < 0.001) with respect to control wild type plants (Fig. 5B). The effect on root development was also evident during adventitious roots formation in *in-vitro* plants (Supplementary Fig. S3A and B). Indeed, root primordia emerged earlier in *35S:CTG134* scions than in wild type, although the root growth was slower, resulting at the end in shorter roots (Supplementary Fig. S3C). Within the hypothesis of the auxin-ethylene crosstalk, the putative mediating role of CTG134 was investigated exposing 35S:CTG134 transformed tobacco plants to ethylene (10μL L^−1^) and grown in dark. As revealed by Environmental Scanning Electron Microscopy (ESEM, Figures 5c–f), although the difference in the root hair phenotype was confirmed, a clear distinction between transgenic lines and controls for the apical hook and hypocotyl thickening, typical of the triple ethylene response, was not observed. Indeed, the untreated (air) 35S:CTG134 (Fig. 5E) seedlings displayed a phenotype similar to those grown in presence of ethylene (Fig. 5D), despite the fact that samples were partially dehydrated by the light vacuum imposed during the ESEM observation. Interestingly, the ethylene treatment induced an additional phenotype in the 35S:CTG134 lines, provoking the development of a massive root hair formation, completely wrapping the root body (Fig. 5F). Subsequently, a Scanning Electron Microscopy (SEM) analysis disclosed that the previously observed root hair phenotype was due to an increase of their density in the 35S:CTG134 lines (Fig. 5H) with regards to control (Fig. 5G). Indeed, most of the root epidermal cells of 35S:CTG134 seedlings developed root hairs, while in WT trichoblasts were arranged in alternating files with atrichoblasts along the root surface.

**Fig 5.**
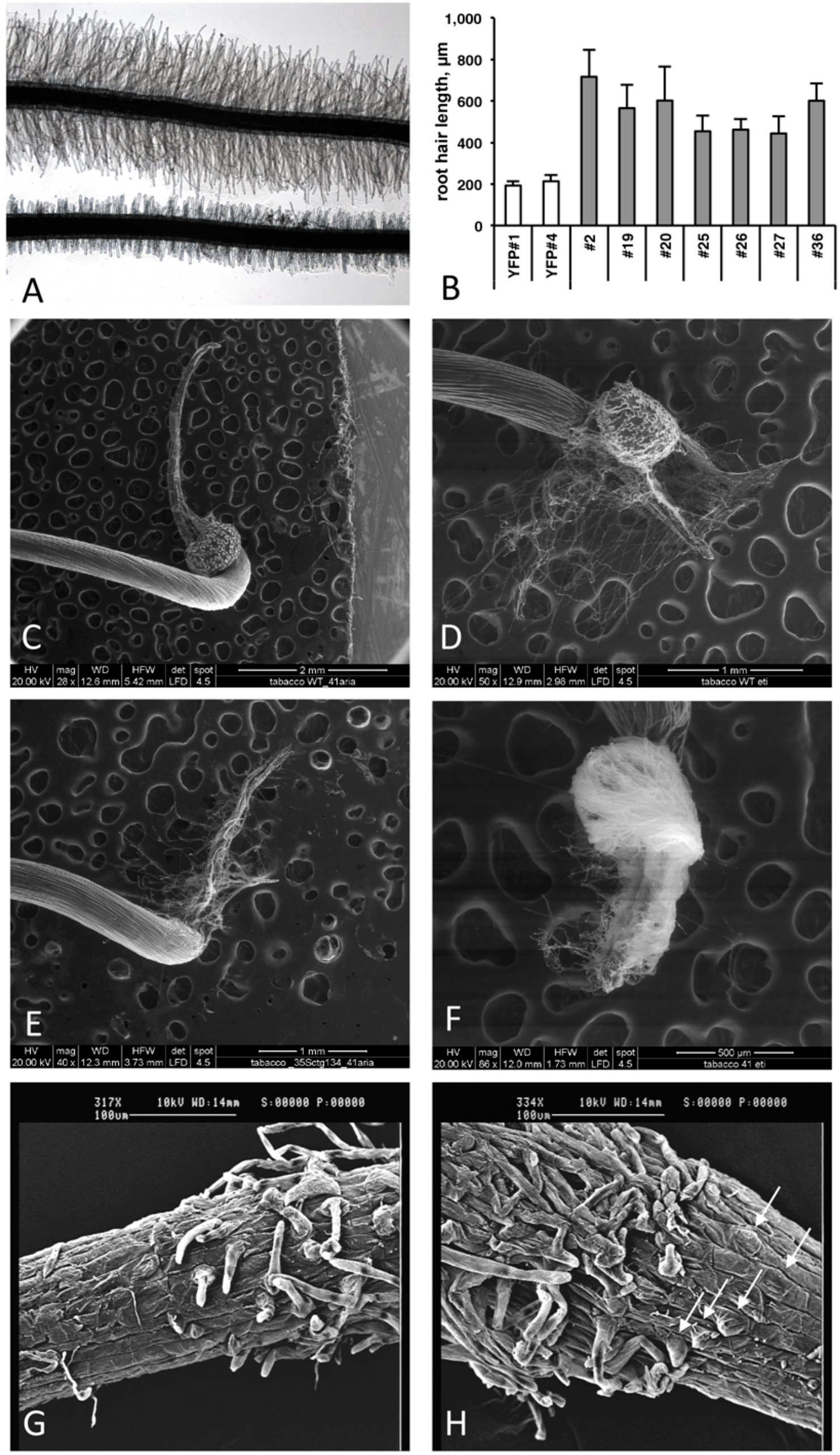
Effects on root growth of *CTG134* overexpression in tobacco. (A, B) CTG134 increases hair length (ANOVA, F = 87.75, df = 155, p < 0.001) in tobacco plantlets grown on agar (controls are transgenic plants expressing the YFP reporter, on bottom in panel A). *CTG134* overexpression in tobacco did not saturate ethylene effect on root hair development and changed the developmental fate of epidermal cells. WT (C, E and G) and *35S:CTG134* (D, F and H) seedling roots were imaged by ESEM after growth in air (C and E) or ethylene (D and F). SEM images of the transition zones of tobacco etiolated seedling roots grown in air showed trichoblasts and atrichoblasts in the WT (G) while almost all epidermal cells were trichoblasts in *35S:CTG134* plants (H; white arrows indicate the presence of root hair primordia that are emerging from epidermal cells).

Since the CTG134 sequence was originally isolated from peach fruit, and placenta cells were stained in tobacco plants expressing the GUS reporter gene driven by the CTG134 promoter, tobacco transgenic capsules were also analysed. Even if tobacco produces a dry fruit structurally different from the fleshy stone fruit of peach, the CTG134 overexpression led to a detectable effect. Tobacco capsules of 35S:CTG134, harvested 12 days after anthesis (before drying), showed an increase in diameter of about 16% with respect to wild type or 35S:YFP (ANOVA, F = 3,85, df = 22, p = 0.013) (Supplementary Fig. S4).

### *Over-expression of* CTG134 *in Arabidopsis*

Similarly to tobacco, the same construct was further employed to transform *Arabidopsis.* T2 CTG134 overexpressing lines were easily identified for their root phenotype when grown on horizontal plates. The primary root of five-day-old 35S:CTG134 seedlings had indeed longer hairs than WT ones (Fig. 6A). Moreover, root hairs developed closer to the apex that in WT roots. To quantify the latter effect, the hairless portion of the root was about half (ANOVA, F = 101.1, df = 23, p < 0.001) of that in the WT (Fig. 6B). As regards to root hair length, being not uniform along the root and clearly depending on age, sizes were taken at given distances from the root-stem transition zone and in a region of the tip that was determined to be, based on growth rate, four-day old. Both measures clearly indicated that the root hairs in the overexpressing lines were longer (ANOVA, F = 95.07, df = 342, p < 0.001; ANOVA, F = 98.31, df = 342, p < 0.001, respectively) than wild type (Fig. 6C). Members of the RGF/GLV family in *Arabidopsis* are known to induce developmental defects in roots when over-expressing seedlings were grown on tilted plates, as reported by (Whitford *et al*., 2012; Fernandez *et al*., 2013). Accordingly, in this work *Arabidopsis* 35S:CTG134 seedlings produced roots with larger and more irregular waves than the WT (Fig. 6D). This effect could be phenocopied by the WT when the synthetic CTG134 peptide (pCGT134) was added to the medium, with the sulfated form being more active than the non-sulfated one (Fig. 6D). Albeit the hairless portion of the root was shorter in overexpressing seedlings, the meristematic region of the root was longer. Moreover, both 35S:CTG134 lines and WT seedlings grown in a medium supplemented with pCTG134 had an increase in root meristem size (Figures 6E and F). The effect on the root meristem size was saturable, as overexpressing lines did not respond to exogenous pCTG134 as the WT (Fig. 6F).

**Fig 6.**
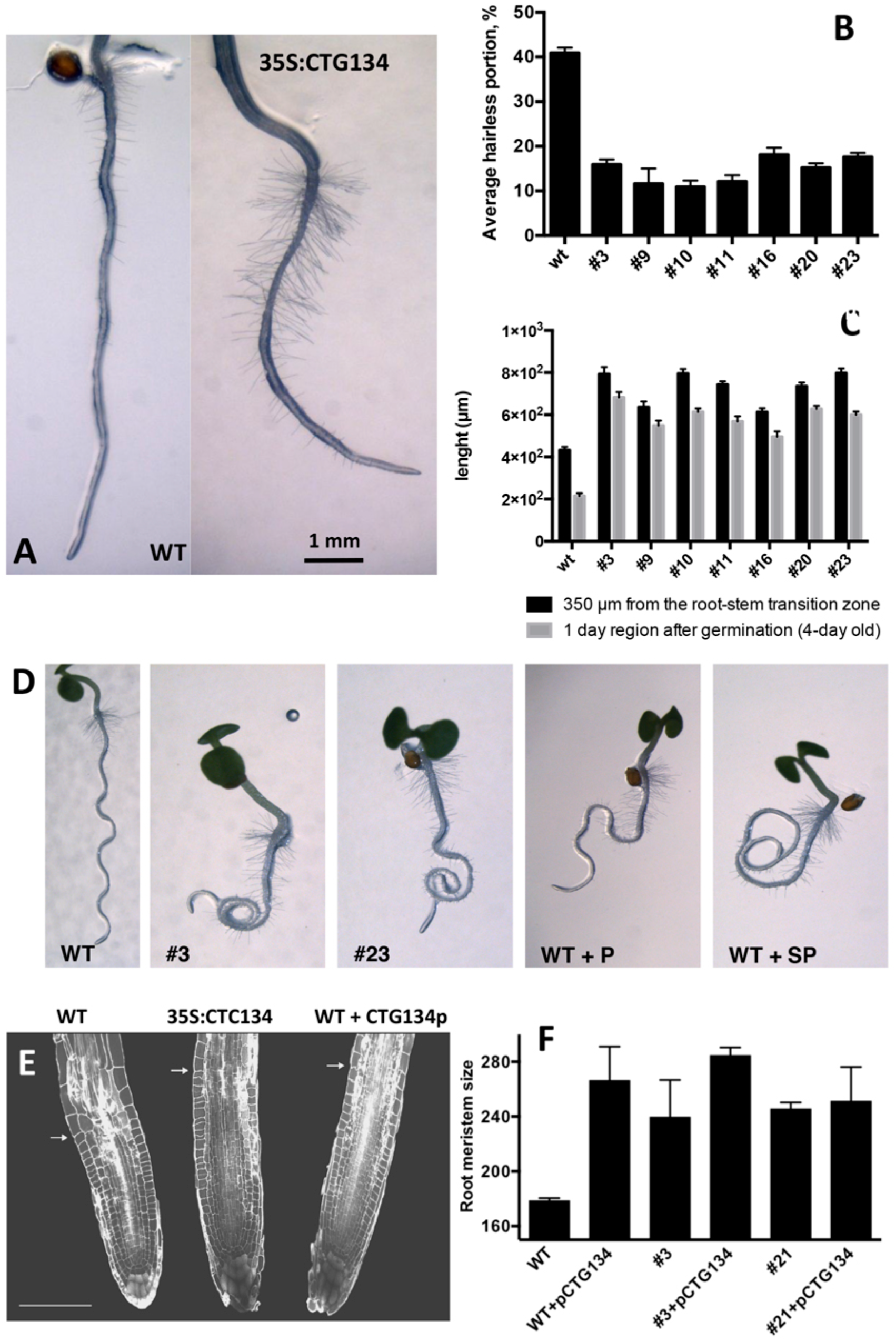
Effects on root development of *CTG134* overexpression in *Arabidopsis.* CTG134 increased hair length (Measurements at 350μm from the root-stem transition zone: ANOVA, F = 95.07, df = 342, p < 0.001; Measurements at 1 day region after germination: ANOVA, F = 98.31, df = 342, p < 0.001) in *Arabidopsis* plantlets grown on agar (A and C); moreover, the portion without root hairs was reduced (ANOVA, F = 101.1, df = 23, p < 0.001) (A and B). Effects on root gravity perception of CTG134 in *Arabidopsis* (D) WT seedlings grown on oblique agar plates showed roots with a regular wavy patter that was altered in CTG134 overexpressing lines (D, #3 and #23). Alteration of the wavy pattern was observed also on WT seedlings grown with synthetic CTG134 peptide added to the medium (D). The effect was stronger if the added peptide was tyrosine-sulfated (WT+SP) compared to the non-sulfated form (WT+P). Effects on root meristem size of CTG134 in *Arabidopsis* (E and F). *Arabidopsis* root sections at five DAG, stained with propidium iodide (e: WT, 35S:CTG134 = overexpressing line, WT + CTG134p = WT grown in the presence of a tyrosine-sulfated synthetic CTG134 peptide). White arrows indicate the transition zone. Scale bar = 100 μm. Measures of meristem size (F) were statistically (Tukey’s multiple comparisons test) larger in comparisons among WT and overexpressing lines (#3 and #21), WT grown in the presence of a tyrosine-sulfated synthetic CTG134 peptide (WT+pCTG134) and overexpressing lines grown in the presence of a tyrosine-sulfated synthetic CTG134 peptide (#3+pCTG134 and #21+pCTG134). Meristem sizes were not statistically different if WT was excluded. Root meristem was measured using Image J software.

The effect of *CTG134* overexpression at the transcriptional level was tested on five-day-old seedling roots (Fig. 7). Alteration in root hairs morphology and quantity was accompanied with a reduction of *GLABRA2* (*GL2*) and a slight induction of *CAPRICE* (*CPC*) expression. The increased meristem size was supported by the expression of *CYCLIN B1;1* (*CYCB1;1*). The development of root hair was selected as a suitable developmental process to test the effect of CTG134 on the interactions between ethylene and auxin occurring at the onset of peach ripening, since the crosstalk of the two hormones during root hair development is well documented (reviewed by Poel *et al.* 2015). The expression of the ethylene biosynthetic gene *ACS2* was induced in roots of 35S:CTG134 seedlings (Fig. 7), as well as that of *ETR1* and *EIN3*, encoding an ethylene receptor and a transcription factors starting the transcriptional cascade leading to ethylene responses, respectively. On the contrary, transcription of *CTR1*, encoding the first downstream signalling component after the ethylene receptor(s) (Kieber *et al*., 1993) was unaffected (Supplementary Fig. S5). About auxin, both *TAA1* and *YUC3* and *6* genes involved in the indole-3-pyruvic acid branch of the hormone synthesis pathway (Tivendale *et al*., 2014) were induced in *CTG134* overexpressing seedlings, while *AMI1*, involved in the indole-3-acetamide branch of the pathway, seemed unaffected (Figures 7 and S5). Free auxin levels depend not only on hormone synthesis but also on its release from storage compartments and transport. The expression of *lAR3*, a gene encoding an IAA-Ala hydrolase (Davies *et al*., 1999), decreased in *CTG134* overexpressing plants, while *PIN2*, encoding an auxin efflux carrier (Müller *et al*., 1998) was induced (Fig. 7).

**Fig 7.**
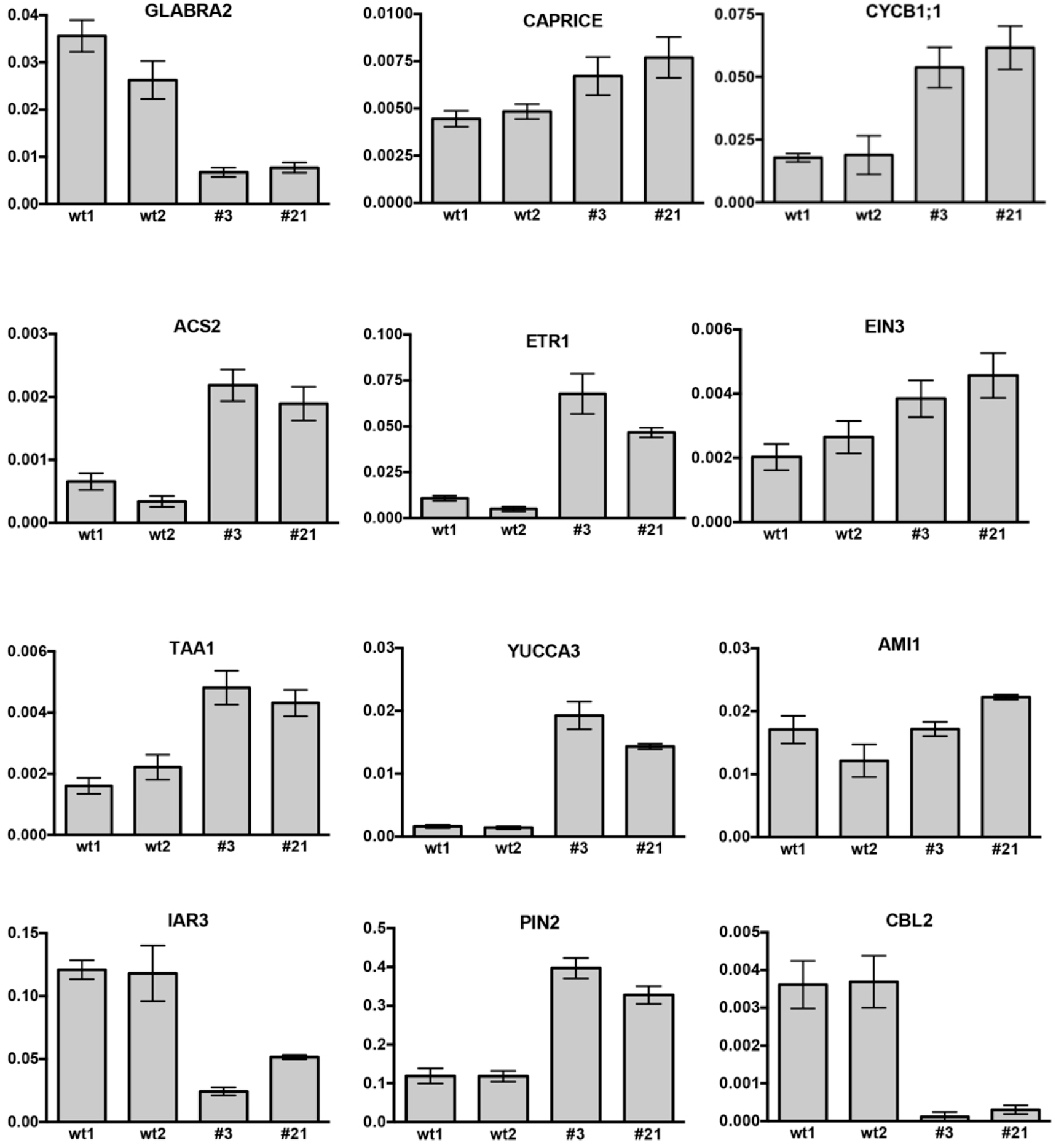
Relative expression profiles of selected genes in roots of *Arabidopsis* seedlings grown on agar plates for five days. wt1 and wt2 are wild type samples collected from two different plates, while #3 and #21 are the clone identifiers of the *Arabidopsis* lines overexpressing the peach *CTG134* gene. Values (means of the normalized expression) have been obtained by real-time qRT-PCR analyses. Bars are the standard deviations from the means of three replicates. ACT8 was used as reference gene.

### pCTG134 induces a cytosolic Ca^2+^ change

In peach, a gene encoding a Ca^2+^ sensing protein belonging to the Calcineurin B-like (CBL) family (*CTG85*) mirrored the expression of *CTG134* during fruit ripening, as well as after 1-MCP treatment (Tadiello *et al*., 2016). The expression of *CBL1, 2, 4*, and *10* encoding genes was therefore tested in 35S:CTG134 roots, showing a general repression, with *CBL2* as the most severely down-regulated gene (Figures 7 and S5).

Given the effect on CBL gene expression and the potential involvement of Ca^2+^ in the signalling pathway activated by signalling peptides (Ma *et al*., 2013), *Arabidopsis* cell suspension cultures stably expressing the bioluminescent Ca^2+^ reporter aequorin in the cytosol were challenged with 100 μM pCTG134. Ca^2+^ measurement assays demonstrated the induction of a biphasic cytosolic Ca^2+^ transient, characterized by a rapid rise, which equally quickly dissipated, followed by a slower Ca^2+^ increase, peaking at about 0.5 μM after 100 s and falling back to basal levels within 5 min (Fig. 8A). No changes in cytosolic Ca^2+^ concentration ([Ca^2+^] _cyt_) were detected in response to either plant cell culture medium (Fig. 8B) or a non-specific peptide (100 μM T16E S19A2) (Fig. 8C), supporting the specificity of the observed Ca^2+^ response to the sulfated CTG134 peptide.

**Fig 8.**
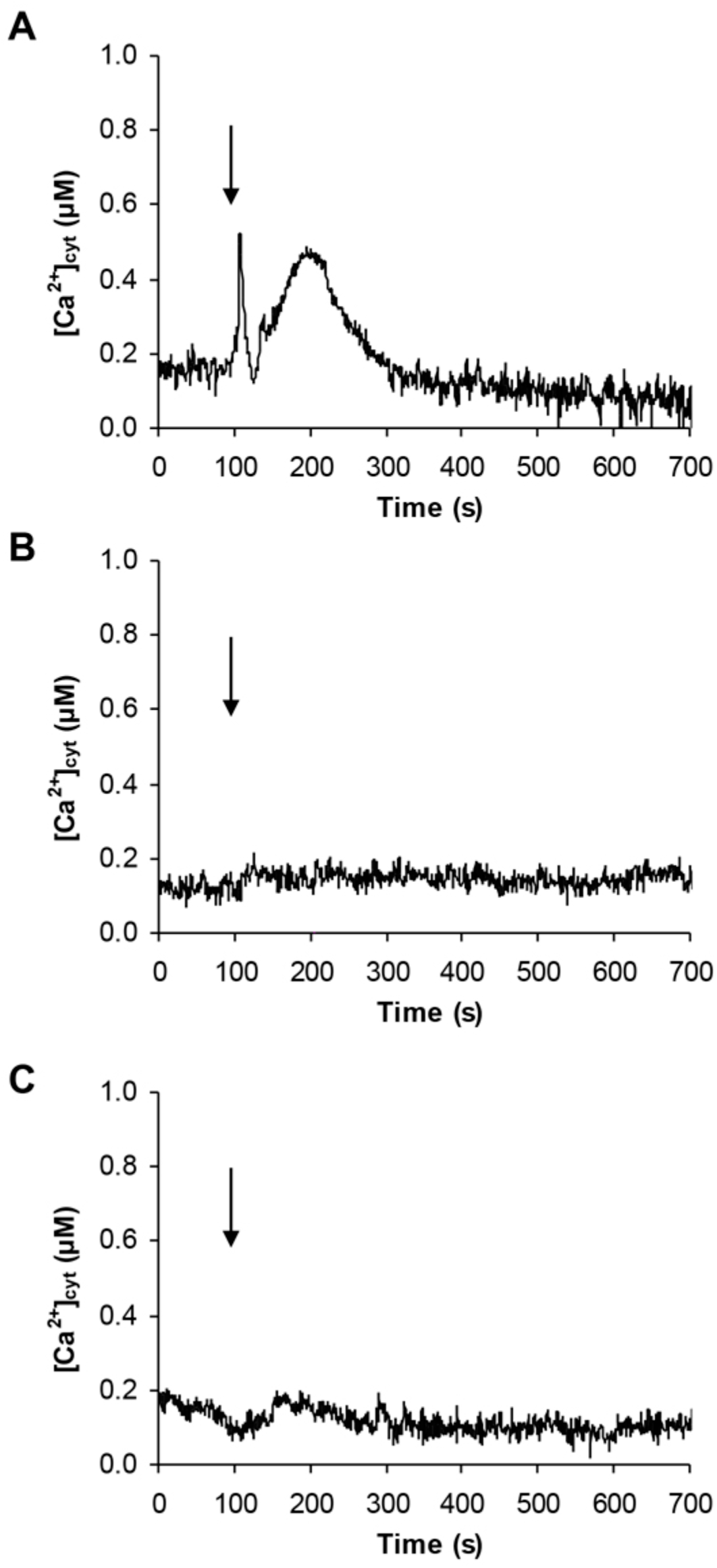
Induction of a transient cytosolic Ca^2+^ change by the sulphonated peptide CTG134S in *Arabidopsis.* Cytosolic Ca^2+^ concentration ([Ca^2+^]cyt) was monitored in aequorin-expressing *Arabidopsis* cell cultures in response to 100 μM CTG134S (A). As controls, cells were challenged with plant cell culture medium (B) or with the non-specific peptide T16E S19A2 (100 μM, C). The arrow indicates the time of injection (100 s). Ca^2+^ traces are representative of three independent experiments which gave very similar results.

## Discussion

Peptide hormones participate in both proximal and distal cell-to-cell communication processes necessary during growth as well as to cope with biotic and abiotic stimuli (reviewed in (Matsubayashi, 2014; Tavormina *et al*., 2015; Wang *et al*., 2016). Despite the growing interest in peptide hormones, their possible role during fleshy fruit ripening remains almost unexplored (Zhang *et al*., 2014). In peach fruit, gene expression profiling suggested that *CTG134*, encoding a peptide belonging to the RGF/GLV family, could be involved in the crosstalk between auxin and ethylene occurring at the onset of fruit ripening (Tadiello *et al*., 2016).

### CTG134 expression is ripening specific and affected by auxin and ethylene perception

Extensive RNA profiling confirmed that *CTG134* is expressed almost exclusively at the onset of ripening, during the transition stage from system 1 to 2 (Fig. 1), as initially suggested by Tadiello *et al.* (2016).

Considering the difficulties typical of *Prunus* species during the *in vitro* regeneration phase, tobacco and *Arabidopsis* transgenic lines expressing the GUS reporter gene driven by the CTG134 promoter sequence, were created. The cis-regulatory elements present in the peach *CTG134* promoter drive *GUS* gene expression in cell/tissue types where the crosstalk between auxin and ethylene was described both in tobacco (Fig. 2) and *Arabidopsis* (Fig. 3). These comprise both cells undergoing separation processes, like abscission, dehiscence zones, lateral root primordia (Roberts *et al*., 2002; Kumpf *et al*., 2013), cambium associated cells (Love *et al*., 2009; Sanchez *et al*., 2012) and placenta cells (De Martinis and Mariani, 1999; Pattison *et al*., 2015). The specificity of the GUS staining pattern obtained in heterologous systems was validated by *in-situ* hybridization in peach mesocarp, where *CTG134* expression was more abundant in bundle associated cells (Fig. 1d). It is noteworthy that also regulatory regions of tomato (Blume and Grierson, 1997), apple (Atkinson *et al*., 1998) and peach (Moon and Callahan, 2004) *ACO* genes drove GUS expression more abundantly in bundle than parenchyma cells of tomato pericarp. Besides spatial regulation, also hormone responsiveness within *CTG134* regulatory regions supported the role in the crosstalk between auxin and ethylene (Fig. 1 and 4). Indeed, both on ripening mesocarp and tobacco seedlings, not only IAA had an inductive effect, probably due to the presence of AREs, but also the altered perception of ethylene (due to 1-MCP treatment) stimulated both *CTG134* transcription in ripening fruit and *GUS* accumulation in tobacco seedlings. In ripening peaches 1-MCP induced auxin synthesis (Tadiello *et al*., 2016), and this might be the reason of the *CTG134* induction. 1-MCP treatment might have induced IAA synthesis, and thus GUS expression, also in tobacco seedlings. In roots of *Arabidopsis* treated with silver (also blocking the perception of ethylene; Negi *et al.* 2008) the exogenous application of 1-MCP might have altered the distribution of IAA, leading to *GUS* induction.

### 35S:CTG134 plants show phenotypes related to auxin and ethylene action

When *CTG134* was permanently overexpressed in tobacco and *Arabidopsis* plants (Figures 5 and 6), the most striking effect was related to the length and number of root hairs, mimicking the effect of exogenous treatments with auxin or ethylene (Pitts *et al*., 1998). Adventitious root formation and elongation in tobacco were also affected, as well as capsule size, further supporting the interplay between auxin and ethylene actions. Besides the well-known effect on root hair number and morphology reported for RGF/GLV/CLEL (Whitford *et al*., 2012; Fernandez *et al*., 2013) and CLE peptides (Fiers *et al*., 2005), CTG134 had an impact also on tobacco capsule size. In fact, at maturity, tobacco capsules were 16% larger than WT on average, similarly to carnation flowers treated with ethylene (Nichols, 1976). Ethylene synthesis is necessary for normal ovule development which impacts flower size (De Martinis and Mariani, 1999). The GUS staining in tobacco placenta and the larger capsules in CTG134 overexpressing plants allow therefore to hypothesize that CTG134 may corroborate auxin inductive and ethylene repressive actions during fruit setting (Martínez *et al*., 2013; Shinozaki *et al*., 2015).

### Molecular targets of CTG134 and its role as mediator in the auxin/ethylene crosstalk

The *Arabidopsis* root model was moreover exploited to gain insights into the regulatory circuit associating CTG134 with auxin and ethylene (Figures 6 and 7). The wavy root phenotype and the increase in meristem size were observed in both overexpressing and peptide treated seedlings, confirming previous findings (Matsuzaki *et al*., 2010; Whitford *et al*., 2012). The observed increase in the meristem size was also supported by the induced expression of *CYCB1;1* (Fig. 7), while the down-regulation of *GL2* was in agreement with its repressing role in root hair development (Ishida *et al*., 2008). More interestingly, genes of both auxin and ethylene synthesis, transport and transduction pathways were upregulated in CTG134 overexpressing roots, assigning to this RGF/GLV peptide a role in the auxin/ethylene crosstalk (Stepanova *et al*., 2007). Although we did not carry out a detailed analysis on the effects caused by the local application of CTG134 peptide (that in Arabidopsis controlled the PIN2 abundance in the root meristem by a post-transcriptional mechanism, thus guiding auxin distribution; Whitford *et al*., 2012), we showed that the heterologous overexpression of the peach CTG134 peptide could be sensed in the portion of the root where receptors initiate the signalling cascade (Shinohara *et al*., 2016; Ou *et al*., 2016; Song *et al*., 2016). As for Peps signalling in *Arabidopsis* (Ma *et al*., 2013), aequorin-based Ca^2+^ measurement assays (Fig. 8) demonstrated the induction by the sulfated peptide CTG134 of a remarkable cytosolic Ca^2+^ change, suggesting the likely involvement of Ca^2+^ as intracellular messenger in the transduction pathway activated by this signal peptide. The role of Ca^2+^ is supported also by the downregulation of several CALCINEURIN B-LIKE PROTEIN (CBL) genes in roots of CTG134 overexpressing seedlings, in agreement with the downregulation of a CBL gene in 1-MCP-treated peaches (Tadiello *et al*., 2016). Sensing the peptide also induced the transcription of key genes of ethylene and auxin biosynthesis pathways and thus, reasonably, the levels of these two hormones, which eventually led to the observed phenotypes. While the response in the ethylene pathway is somewhat straightforward investigating the induction of key genes in its synthesis (ACS2), perception (*ETR1*) and signal transduction (*EIN3*), the action on the auxin pathway is more intricate. Indeed, while the increased transcription of *TAA1, YUC3* and *YUC6* sustains the induction of the two-step IPA pathway, the unchanged levels of *AMI1* seemed to exclude the conversion of indole-3-acetamide (IAM) to IAA (Enders and Strader, 2015). Moreover, although only *IAR3* was tested, the contribution of conjugated forms of IAA (Sanchez Carranza *et al.*, 2016) seemed negligible in *Arabidopsis*, while the expression of its peach homolog *CTG475* was supposed to participate to the free auxin increase measured before the climacteric production of ethylene in peach (Tadiello *et al*., 2016), thus complementing the role of *PpYUC11* (Pan *et al*., 2015). However, the induced transcription of *PIN* genes in overexpressing *Arabidopsis* seedlings (Fig. 7) and in climacteric peaches (Tadiello *et al*., 2016) supported a key role of these peptides in regulating auxin distribution (Whitford *et al*., 2012).

The comprehensive expression profiling data carried out in peach (Tadiello *et al*., 2016) and the knowledge here achieved about CTG134 in tobacco and *Arabidopsis* provide evidence on the involvement of this RGF/GVL secreted peptide in a regulatory circuit that sustains auxin and ethylene actions. The same circuit, working in both rosids *(Arabidopsis)* and asterids (tobacco) might have appeared early during evolution of eudicots to participate in the control of root hair development and later it could have been recruited in peach to regulate the switch from system 1 to system 2 ethylene synthesis (Fig. 9). Further research will be necessary to clarify the molecular details by which CTG134 acts to either regulate auxin and ethylene synthesis or modify their distribution and perception, or both. The kinase nature of GLVs receptors (Shinohara *et al*., 2016; Ou *et al*., 2016; Song *et al*., 2016) agrees with the measured Ca^2+^ perturbations.

**Fig 9.**
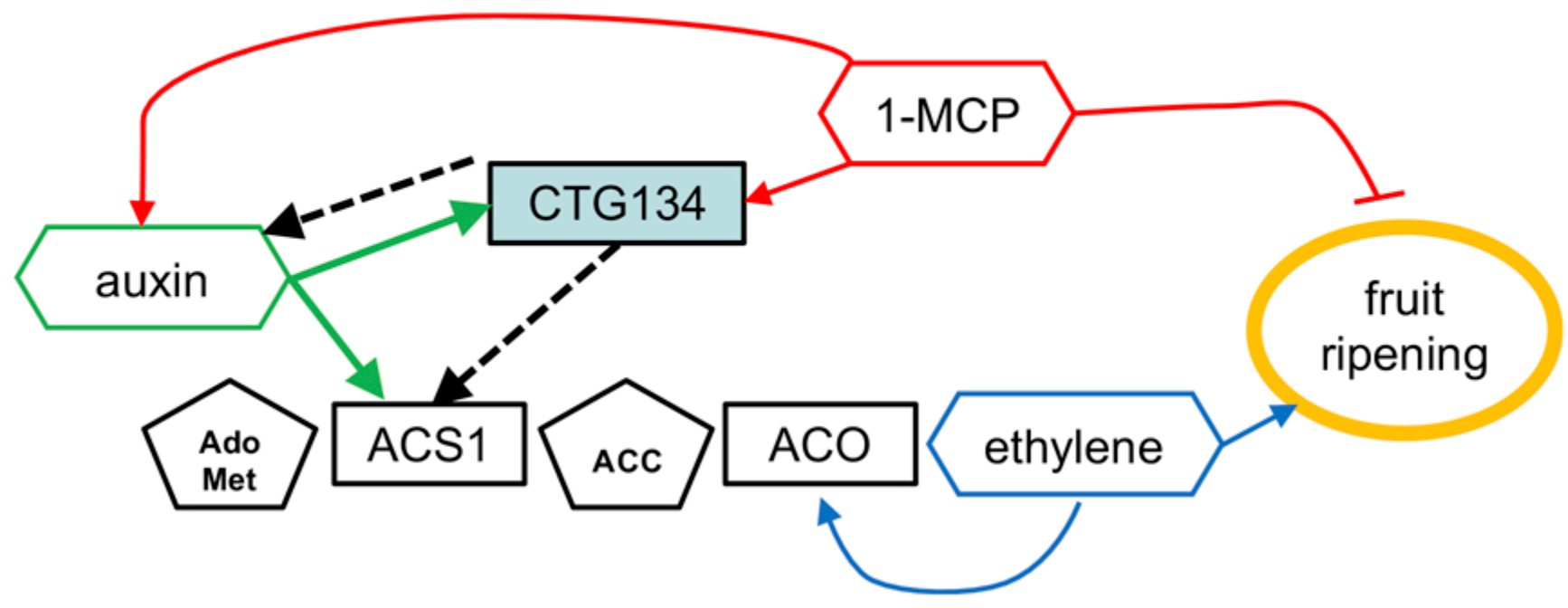
A model positioning *CTG134* in the regulatory network controlling peach ripening. Regulatory data collected from the *Arabidopsis* CTG134 overexpressing clones are represented by dashed lines. Ethylene autocatalytic synthesis and action on fruit ripening is represented in blue, auxin, 1-MCP and CTG134 interactions in green, red and black, respectively. Hormones (or inhibitors) are in hexagons, their precursors in pentagons while genes (gene products) are in rectangles. Filled arrow means induction, while blunted lines repression.

The unique mechanism that switches ethylene synthesis from system 1 to system 2 in peach probably relies on the use of a single ACS gene for both kinds of syntheses (Tadiello *et al*., 2016), thus differing from tomato (Barry *et al*., 2000) and apple (Wang *et al*., 2009). In these two latter fruits, the expression of *LeACS4* and *MdACS3* (system 1) is necessary to start *LeACS2* and *MdACS1* transcription (system 2), respectively. During peach ripening, expression of other ACS genes is, if present, several orders of magnitude lower than that of *ACS1* (Tadiello *et al*., 2016). The different amount of ethylene released by system 1 and system 2 could be achieved by modulating system 1 ACS1 activity, thus leading to system 2 *ACS1* increased transcription. ACS1 belongs to type-1 ACS proteins, which are stabilized by phosphorylation mediated by mitogen-activated protein kinases (MAPKs) (Liu and Zhang, 2004). Phosphorylation cascades have been shown to start upon binding of peptide signals (e.g. IDA) with their receptors (e.g. HAE/HSL2) (Cho *et al*., 2008). Given the transcriptional regulation of *CTG134*, the nature of pCTG134 and of the *Arabidopsis* receptors of its homologous RGF/GLV peptides (Shinohara *et al*., 2016; Ou *et al*., 2016; Song *et al*., 2016) and of the ability of pCTG134 to trigger a cytosolic Ca^2+^ signal, we hypothesized that the transition of ethylene synthesis from system 1 to system 2 in peach could be controlled by ACS1, whose activity might be therefore modulated through the action of pCTG134.

## Supplementary Data

**Fig. S1**. Details of the vectors used for gene overexpression and promoter analyses.

**Fig. S2**. Localization of *CTG134* expression in peach mesocarp by *in-situ* hybridization (control panels).

**Fig. S3**. Adventitious root formation in 35S:CTG134 clones and in control lines.

**Fig. S4**. Effects on capsule size of *CTG134* overexpression in tobacco.

**Fig. S5** Relative expression profiles of selected genes in roots of *Arabidopsis* seedlings grown on agar plates for five days.

**Table S1** List and sequences of DNA primers used.

## Acknowledgements

We thank M.R. Knight (Durham, UK) and G. Regiroli (AgroFresh Inc., Philadelphia, PA, USA) for kindly providing seeds of aequorin-expressing Arabidopsis plants and SmartFreshTM (1-MCP), respectively. The authors are also grateful to Alice Tadiello and Maria Patrizia Schiappelli for providing preliminary expression data and support in peptide synthesis. Financial support was provided by MIUR (Italian Ministry of Research and University), MiPAFF (Ministero delle Politiche Agricole Alimentari e Forestali–Italy; www.politicheagricole.it) through the project ‘DRUPOMICS’ (grant DM14999/7303/08) and the University of Padova (grant CPDA072133/07 and CPDA132841/13) to LT.

